# GFP Reporter System Reveals Cell-to-Cell Variability in Aquaporin-2 Expression

**DOI:** 10.64898/2025.12.16.694664

**Authors:** Lihe Chen, Adrian Murillo-de-Ozores, Euijung Park, Shuo-Ming Ou, Mark A. Knepper

**Author notes:** Correspondence: Mark Knepper, MD, PhD, Division of Intramural Research, National Heart, Lung and Blood Institute, NIH, Bethesda, MD 20892.

## Abstract

Vasopressin regulates transcription of the aquaporin-2 gene (*Aqp2*) in collecting duct principal cells. To investigate regulatory mechanisms in *Aqp2* gene transcription, we engineered an *Aqp2* reporter cell line using CRISPR/Cas9 to insert a green fluorescent protein (GFP) cassette at the endogenous *Aqp2* gene locus in mpkCCD cells. In the absence of dDAVP, a vasopressin analog, these cells exhibited low or undetectable GFP and Aqp2 expression in all cells. dDAVP stimulation (1nM dDAVP for 48hrs) markedly increased both GFP and Aqp2 expression together with reversal upon dDAVP removal. These observations demonstrate that GFP faithfully tracks Aqp2 expression. Interestingly, fewer than 50% of cells express GFP and Aqp2 after dDAVP or forskolin, indicating significant variability even though they were clonally-derived. We flow-sorted the GFP^−^ cells (Aqp2^−^) and GFP^+^ cells (Aqp2^+^), regrew them, and restimulated them separately with dDAVP. Cells originating from GFP^−^ cells gave rise to both GFP^−^ cells and GFP^+^ cells, and GFP^+^ cells similarly regenerated both GFP^−^ and GFP^+^ populations in the same proportion. Flow cytometry analysis of the DNA content showed variability in cell cycle phases, with most GFP^+^ cells in G0/G1, and more GFP cells in G2/S. RNA-seq analysis of the GFP^−^ and GFP^+^ cells revealed increased abundance of cell-cycle related transcripts in the GFP^−^ cells. We conclude that: 1) heterogeneity in Aqp2 expression is related to cell cycle state; and 2) the newly generated reporter cell line will likely serve as a useful tool to study *Aqp2* transcriptional regulation.

**NEW & NOTEWORTHY:** To investigate regulatory mechanisms in *Aqp2* gene transcription, we engineered an *Aqp2* reporter cell line using CRISPR/Cas9 to insert a green fluorescent protein (GFP) cassette at the endogenous *Aqp2* gene locus in mpkCCD cells. We demonstrate that the GFP reporter accurately and dynamically tracks the expression and regulation of endogenous Aqp2. We reveal that Aqp2 heterogeneity in mpkCCD cells is at least partly driven by differences in cell cycle phase.

## INTRODUCTION

Cell-to-cell heterogeneity is a fundamental phenomenon in cellular systems, where even genetically identical populations can display significant variability in gene expression and responses to stimuli (1). This has been widely recognized in diverse contexts, including developmental plasticity (2, 3), tumor heterogeneity (4, 5), and drug resistance (5, 6). Recent advances in single-cell genomics and advanced microscopy have greatly improved our understanding of cellular variation (7, 8), but the question of how cells with identical DNA sequences give rise to such variability still remains an active area of research.

Cellular variation has been documented within the kidney collecting duct, which plays a critical role in maintaining water, electrolyte, and acid-base balance (9). The collecting duct principal cells are responsible for water reabsorption, a process primarily mediated by the water channel protein, Aquaporin-2 (Aqp2) (10-12). The expression and trafficking of Aqp2 are tightly regulated by the hormone arginine vasopressin (AVP) (9, 10, 13-15). Disruption of this process leads to nephrogenic diabetes insipidus (NDI), a condition characterized by the kidney’s inability to concentrate urine (16-19).

The mouse collecting duct cell line (mpkCCD) is widely used as a cell model to study the actions of vasopressin. These cells recapitulate most of the regulatory functions seen in native kidney collecting duct principal cells (20, 21). While the overall vasopressin-mediated regulation of Aqp2 is well-documented, studies have shown that not all the cells respond uniformly to AVP stimulation (22). The underlying mechanisms behind this cell-to-cell heterogeneity in Aqp2 expression are not fully understood, partially due to lack of an efficient tool to track the Aqp2 changes in live cells.

To provide a surrogate marker for Aqp2 expression and to investigate the mechanisms of vasopressin-mediated heterogeneity, we developed an Aqp2-GFP reporter cell line using CRISPR/Cas9-mediated gene editing—a technique that has revolutionized genetic engineering (23). By tagging the endogenous *Aqp2* gene with GFP in mouse collecting duct cells (mpkCCD), we were able to directly visualize and quantify Aqp2 expression in living cells. Here, we show that GFP faithfully tracks the Aqp2 expression and could serve as a reporter for Aqp2 transcriptional regulation. We further investigate the heterogeneity of Aqp2 expression using RNA-seq and FACS analysis and show that the heterogeneity is partially driven by the cell cycle. Finally, our reporter cell line could provide as a powerful tool for studying Aqp2 transcriptional regulation in future studies screening for both positive and negative regulators.

## MATERIALS AND METHODS

### Cell Culture

Mouse cortical collecting duct cells (mpkCCDc11 secondary clone 38, mpkCCD_C11-38_) were grown on transwells (Cat. No. 3460 and 3450, Corning) in complete DMEM/Ham’s F12 medium (DMEM/F12) containing 5 μg/mL insulin, 50 nM dexamethasone, 1 nM triiodothyronine, 10 ng/mL epidermal growth factor, 60 nM sodium selenite, 5 μg/mL transferrin, and 2% fetal bovine serum for 4 days. Then, the medium was changed to a serum-free/growth factor-free simple medium [with 50 nM dexamethasone, 60 nM sodium selenite, 5 μg/mL transferrin] to ensure complete polarization and cell differentiation. Cells were treated with vehicle or 0.1 nM or 1 nM dDAVP on the basolateral side for 24 or 48 hours in simple medium. To test Forskolin effect on the cells, 10 μM Forskolin were added on the both sides for 48 hours in simple medium.

### Generation of Aqp2 reporter cell lines

We engineered the *Aqp2* reporter cell lines by inserting a green fluorescent protein (GFP) cassette at the *Aqp2* gene locus in mpkCCD cells using the CRISPR/Cas9. First, we designed a specific sgRNA (5’ CGAGCAGCATGTGGGAACTC 3’) targeting the initial codon (ATG) of the *Aqp2* gene using the CHOPCHOP application (https://chopchop.cbu.uib.no/). The sgRNA is then cloned to a vector harboring the Cas9 and a red fluorescent protein RFP sequence (Vector No. p02, sigma). Next, a donor DNA template was synthesized by GenScript (https://www.genscript.com/) which contains a GFP sequence and additional homologous sequences flanking 3 kb 5’ and 2.5 kb 3’ of *Aqp2* (See Supplemental Fig. S1 for sequence). A P2A cleavage sequence was introduced between the GFP and the open reading frame (ORF) of Aqp2. EcoRI and SalI restriction sites flanking both the end of the donor fragment was then cloned into pUC57. Finally, the sgRNA plasmid (sgRNA + Cas9 + RFP) and the donor plasmid (5’ flanking sequence + GFP + P2A + 3’ flanking sequence) were transfected into mpkCCD cells using lipofectamine 3000 (L3000001) according to the manufacturer’s instruction. 48 hours after transfection, the cells were sorted into transwell based on RFP expression. Cells with potential edits were cultured in complete medium with 0.1 nM dDAVP for 7 days followed by single-cell GFP sorting. More than 50 potential single *Aqp2*:GFP reporter clones were obtained and the final clones were characterized and selected based on, 1) Primer set (5’ GGTCTACGATAGGAAGGCCC 3’ and 5’ GAAGACGAAAAGGAGCGTGG 3’) is designed to target both 5’ flanking region and 3’ flanking region to identify the WT allele (200 bp) and GFP inserted allele (1000 kb); 2) Primer set (5’ GCAGATACGAGCCACACTTTC 3’ and 5’ tccttgaagaagatggtgcg 3’) and primer set (5’ aagttcatctgcaccaccg 3’ and 5’ GCAGCTGGAAGCTTTGAACC 3’) targeting either the upstream region of the flanking sequence or the downstream region of the flanking sequence together with GFP to identify on target GFP insertion; 3) Sanger sequencing was used to confirm the presence of the GFP sequence at the intended site.

### Immunoblotting

Cells were rinsed twice with ice-cold Dulbecco’s phosphate-buffered saline (DPBS) and then lysed with lysis buffer (1.5% SDS, 10 mM Tris, pH 6.8) containing protease and phosphatase inhibitor (78441, Thermo Fisher Scientific). The lysates were collected and passed through the QIAshredder (79656, QIAGEN). Samples were denatured and subjected to SDS-PAGE using 4-20% or 12% Criterion TGX Gels (5671095, 5671045, Bio-Rad) and then transferred to nitrocellulose membranes. The blots were blocked and incubated with primary antibodies overnight at 4 degree. Primary antibodies were anti-AQP2 (K5007, 1:5000) and anti-GFP (2555S, CST, 1:1000). Fluorescence images were scanned by the Li-COR Odyssey System (ODY-0428). The analyses were done using Li-COR Image Studio software.

### Immunofluorescence Microscopy

Cells were washed and then fixed with 4% paraformaldehyde for 10 min at RT. Cells were washed three times and then blocked with blocking solution (1% BSA and 0.2% gelatin in PBS) for 30 min. The cells were washed again and permeabilized with permeabilizing solution (0.3% Triton X-100 and 0.1% BSA in PBS) for 10 min. Cells were washed and incubated with primary antibodies overnight at 4°C. Primary antibodies was anti-AQP2 (K5007, 1:500). GFP was directly visualized under the microscope. Cell nuclei were stained using DAPI (4⍰,6-diamidino-2-phenylindole). Confocal images were taken using the Zeiss LSM 780 microscope (National Heart, Lung and Blood Institute, Light Microscopy Core Facility). Images were analyzed by *Zen* software.

### Flow cytometry

Cells were washed and dissociated by adding the TrypLE Express Enzyme (12604013, Giboco) to both sizes of the membrane. Samples were collected and passed through the 35 µM cell strainer. The sorting was done at the National Heart, Lung, and Blood Institute (NHLBI) Flow Cytometry Core. Analyses were performed using the *FlowJo* V10 software.

### RNA-seq

mpkCCD cells or reporter clones were grown on transwells and treated with or without dDAVP (1 nM for 48 hours). Total RNA was isolated using Direct-zol RNA Microprep kit (R2062, Zymo Research) following the manufacturer’s instructions. cDNA was generated from 50 ng of total RNA using SMART-Seq v4 Ultra Low Input RNA Kit. 1 ng of cDNA was “tagmented” and barcoded by using a Nextera XT DNA Sample Preparation Kit (Illumina). The final libraries were purified by AmPure XP magnetic beads (Beckman Coulter, Indianapolis, IN) and quantified using a Qubit 2.0 Fluorometer (Thermo Fisher Scientific, Waltham, MA). An equal amount of index libraries were pooled and sequenced (paired-end 50 bp) on an Illumina Novaseq 6000 platform. Raw sequencing reads were aligned by STAR 2.7.10b (24) to the mouse reference genome (Ensembl genome 106). Transcript per million (TPM) and expected counts were generated by RSEM 1.3.1 (25). Differential gene expression analysis were performed using DESeq2 (26). To quantify the GFP mRNA expression level, we created a custom reference by inserting the GFP sequence into the mouse genome.

### Statistics

The enrichment of specific gene ontology biologic processes was done in R using *clusterProfiler* based on Fisher’s Exact Test. Student’s t-test was used to compare the means between two groups including control and dDAVP or FSK treatment in western blotting and FACS experiments.

## RESULTS

### Development of Aqp2-GFP reporter cell line

The mpkCCD cell line, which is well-established and widely used for investigating the transcriptional regulation of Aqp2 (27), was chosen as the parental cell line for generating the Aqp2-GFP reporter using CRISPR/Cas9-mediated gene editing. Our overall strategy is to employ the CRISPR-Cas9 technology to create double strand DNA (dsDNA) breaks in the cells and rely on the homology-directed repair (HDR) mechanism (28) to insert the green fluorescent protein (GFP) coding sequence at the *Aqp2* locus during the DNA repair process. To achieve this, we designed a single guide RNA (sgRNA) spanning the start codon (ATG) of *Aqp2* and cloned it into a vector that also expresses Cas9 and red fluorescent protein (RFP) (Fig. 1A). The RFP served as a marker to select for cells successfully transfected with the CRISPR components. In parallel, we synthesized a DNA repair template (donor) consisting of GFP flanked by 5’ and 3’ *Aqp2* homology arms (Fig. 1A; Supplemental Fig. S1). A P2A sequence was placed between GFP and the Open Reading Frame (ORF) of *Aqp2*, allowing expression of both GFP and full-length Aqp2 proteins from the same transcript.

**Figure 1.**
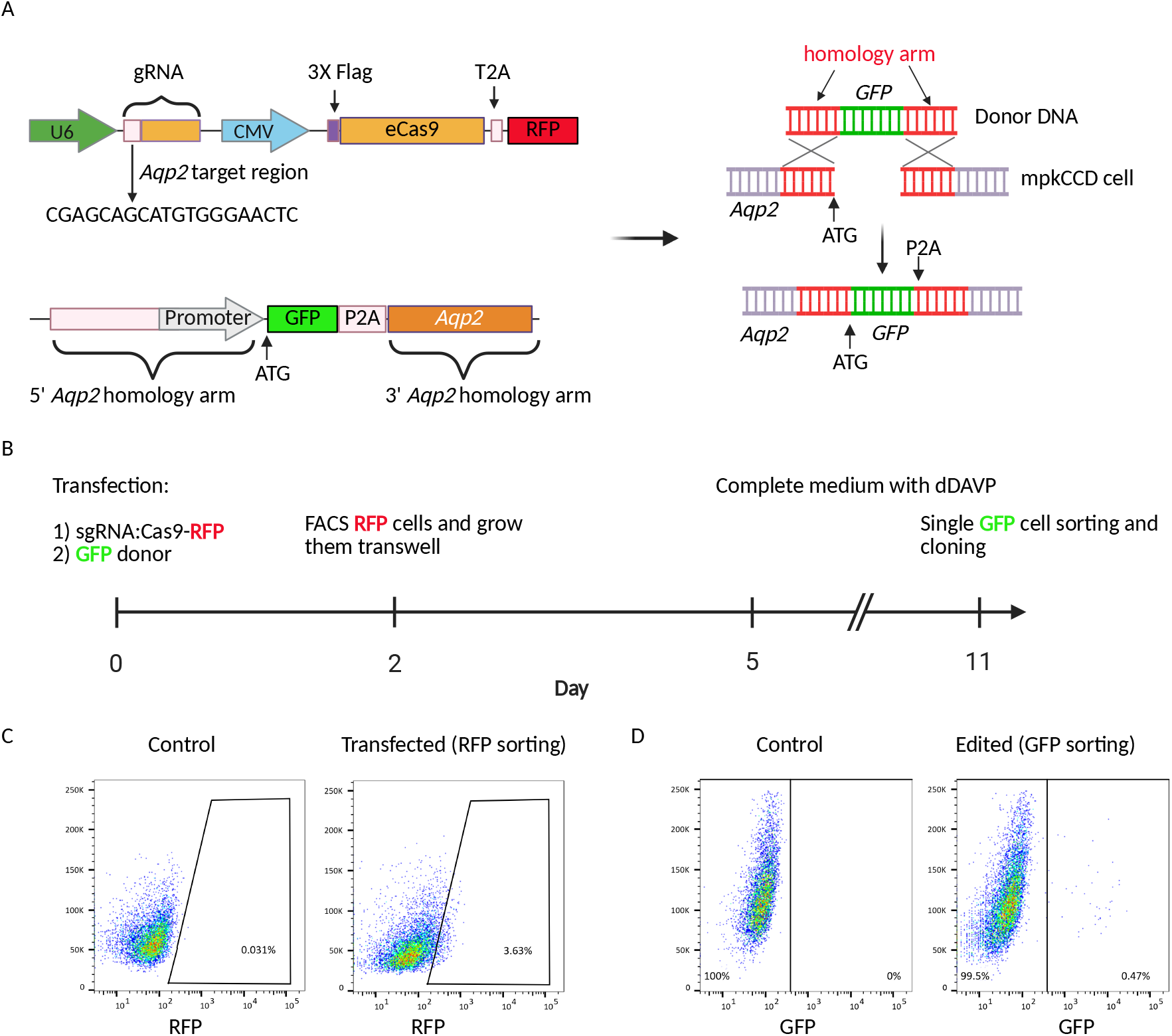
Generation of the Aqp2-GFP reporter cell line. *A*: Two main plasmids are used for the gene editing: the CRISPR vector and the GFP donor DNA. The CRISPR plasmid expresses a sgRNA targeting the initial codon of Aqp2, the Cas9 enzyme, and the RFP. The donor plasmid consists of the GFP sequence, which is flanked by the 5’ and 3’ Aqp2 homology arms. A P2A sequence is placed between GFP and the Aqp2 ORF. Co-transfection of these two plasmids into the mpkCCD cells results in inserting the GFP into the Aqp2 locus through HDR mechanism. *B:* Workflow for generating the Aqp2-GFP reporter cell line. C: FACS sorting of the RFP-positive cells to select those that successfully received the CRISPR plasmid. The plot shows the transfection rate is around 3.6% compared to the control population without transfection. C: FACS sorting of the cells with potential GFP insertion. Only 0.47% of the transfected cells were GFP^+^, which indicates the low but successful rate of HDR.

This specific design offers several key advantages: 1) Insertion of the GFP gene in the vicinity of the *Aqp2* promoter ensures the reporter is under the control of the endogenous Aqp2 regulatory mechanism, thereby guaranteeing GFP accurately tracks Aqp2 expression; 2) Designing a guide RNA (sgRNA) to the start codon prevents the sgRNA from cutting the donor DNA (GFP insertion disrupts the sgRNA recognition site in the donor), which leads to higher HDR efficiency (29); 3) The “self-cleaving” peptides (P2A) (30) separates GFP from Aqp2 during translation, effectively preventing protein fusion.

The plasmids with the CRIPSR components and repair template were co-delivered into the mpkCCD cells (Fig. 1B). The transfected cells (RFP) were subsequently sorted directly into transwell plates by FACS (Fig. 1C) and maintained with dDAVP before final single GFP-positive cell sorting (Fig. 1 D). As expected, the HDR rate is low (31), with only 0.47% of the transfected cells being GFP^+^ (Fig. 1D). Following the expanding the single clones, we obtained over 50 GFP reporter clones and selected a subset for thorough characterization.

### Characterization of Aqp2-GFP reporter cell line

To confirm the precise integration of the GFP reporter gene into the *Aqp2* locus, we used junction PCR (32). As shown in the top panel of Fig. 2A, a primer pair designed with both primers located within the homology arm generated a ∼1000 bp product only when the GFP gene has been successfully integrated. This method confirmed that all tested clones (n=10) were successfully tagged. The simultaneous presence of a ∼ 200 bp wild-type (WT) band confirmed that all clones were heterozygous. To rule out random integration events, we designed a second set of primer pairs as shown in the bottom two panels of Fig. 2A. In this design, one primer is placed within the newly inserted GFP sequence and the other is placed outside the homology arm. This configuration confirms that the GFP gene was correctly inserted at the specific *Aqp2* locus and not elsewhere in the genome. The examples illustrate that 7 out of 10 clones exhibited correct site-specific insertion. Finally, Sanger sequencing was performed on candidate clones to further verify the accuracy of integration and identify possible mutations at the integration site.

**Figure 2.**
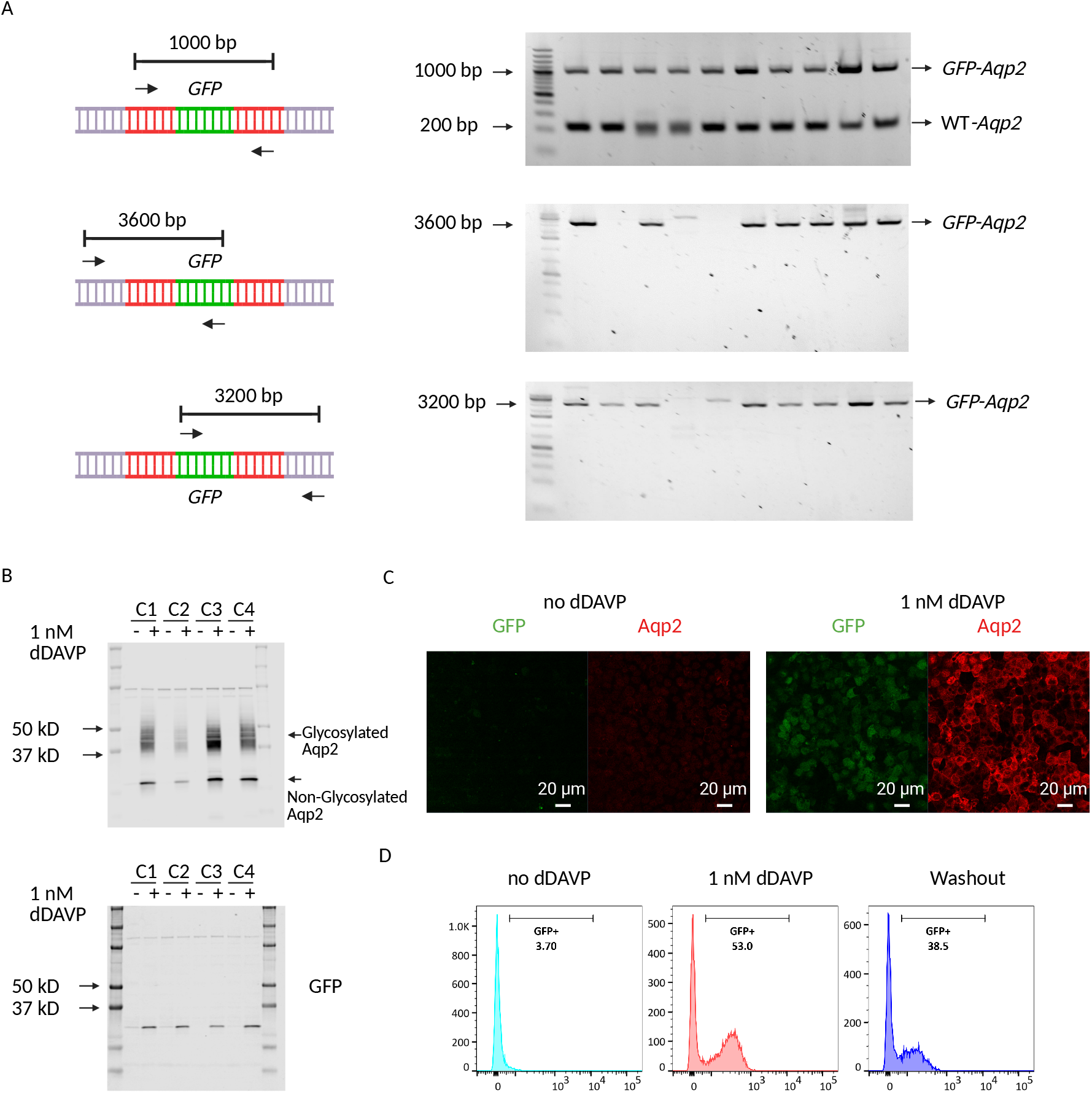
Validation of the Aqp2-GFP reporter cell lines. *A:* Junction PCR strategy to confirm the precise integration of the GFP at the Aqp2 locus. The primer design places both primers within the homology arm of the donor DNA, ensuring a 1000 bp band is generated only upon successful GFP insertion (left top panel). PCR primers are designed with one primer inside of the GFP and the other outside of the homology arm of the Aqp2 (left middle and bottom panels). The design ensures the GFP is inserted at the intended Aqp2 locus. The PCR band sizes are around 3600 bp and 3200 bp respectively. *B*: Semiquantitative immunoblots on four representative clones treated with or without 1nM dDAVP. This confirms that the GFP is regulated similarly to endogenous Aqp2. *C:* Confocal microscopy analysis of the Aqp2-GFP reporter cell line. While GFP and Aqp2 are undetectable in the absence of dDAVP, stimulation (1 nM dDAVP) increases GFP and Aqp2 expression in a subset of cells. *D:* FACS analysis of Aqp2-GFP reporter cell line. The control cells show a low background of GFP signal (3.7%). After addition of dDAVP, the GFP fluorescence shifts dramatically to the right, showing a significant increase in the positive population to 53.0%. Removal of dDAVP decreases the GFP population to 38.5%.

Next, we used semiquantitative immunoblotting, confocal microscopy, and FACS to functionally validate the clones. Clones were grown and differentiated on membrane filters before dDAVP treatment (Methods). As shown in Fig. 2B; Supplemental Fig. S2, semiquantitative immunoblotting showed concurrent expression of Aqp2 and GFP protein following treatment. The absence of both proteins in the untreated samples confirmed the dDAVP-dependent regulation of the GFP reporter, accurately mirroring the endogenous Aqp2 expression. Detection of a single GFP band of the expected size confirmed the efficient cleavage of the GFP from the Aqp2 protein by the P2A peptide.

Colocalization of GFP and Aqp2 was examined by immunofluorescence (Fig. 2C). While expression of both was undetectable without dDAVP, stimulation with dDAVP caused a subset of cells to exhibit increased GFP and Aqp2 expression. Aqp2 maintained its normal physiological function of redistributing to the apical membrane. Although there is an obvious dDAVP-induced expression of GFP and Aqp2, some cells showed no or low expression of both, suggesting a heterogeneous response to dDAVP. This heterogeneity in dDAVP response was corroborated by FACS analysis (Fig. 2D), which showed a significant increase in both the number of GFP-positive cells and their fluorescence intensity following dDAVP treatment. This effect was reversible upon dDAVP removal. These results collectively demonstrate that the GFP reporter accurately and dynamically tracks the expression and regulation of endogenous Aqp2.

### Dynamic expression of Aqp2

To investigate whether the GFP reporter cell line accurately reflects the known time-dependent regulation of Aqp2 by dDAVP (21), we conducted a kinetic analysis. Cells were treated with dDAVP at various timepoints (0, 6, 24, and 48 hours) and subsequently analyzed by FACS and immunoblotting (Methods). As illustrated in Fig. 3A, the proportion of GFP-positive cells markedly increased over time, rising from less than 0.5% to over 3% after 6 hours and exceeding 30% after 24 hours of dDAVP exposure. This increase was accompanied by a corresponding rise in GFP intensity per cell. This finding suggests that the observed increase in total Aqp2 protein expression is likely a cumulative effect of two factors: an increase in the number of Aqp2-expressing cells and an elevated level of Aqp2 expression within each cell. To directly quantify the Aqp2 protein expression, we performed immunoblotting. Fig. 3, B and C show the total Aqp2 protein significantly increased over the dDAVP treatment period. Densitometry analysis (Fig. 3C) demonstrated an increase of more than 250-fold compared to untreated control. This dramatic rise in Aqp2 protein expression directly validates the trend observed with the GFP reporter and confirms that dDAVP treatment leads to a substantial, time-dependent upregulation of Aqp2 protein (21). This result supports the conclusion that the GFP reporter system is a reliable tool for monitoring Aqp2 regulation in these cells.

**Figure 3.**
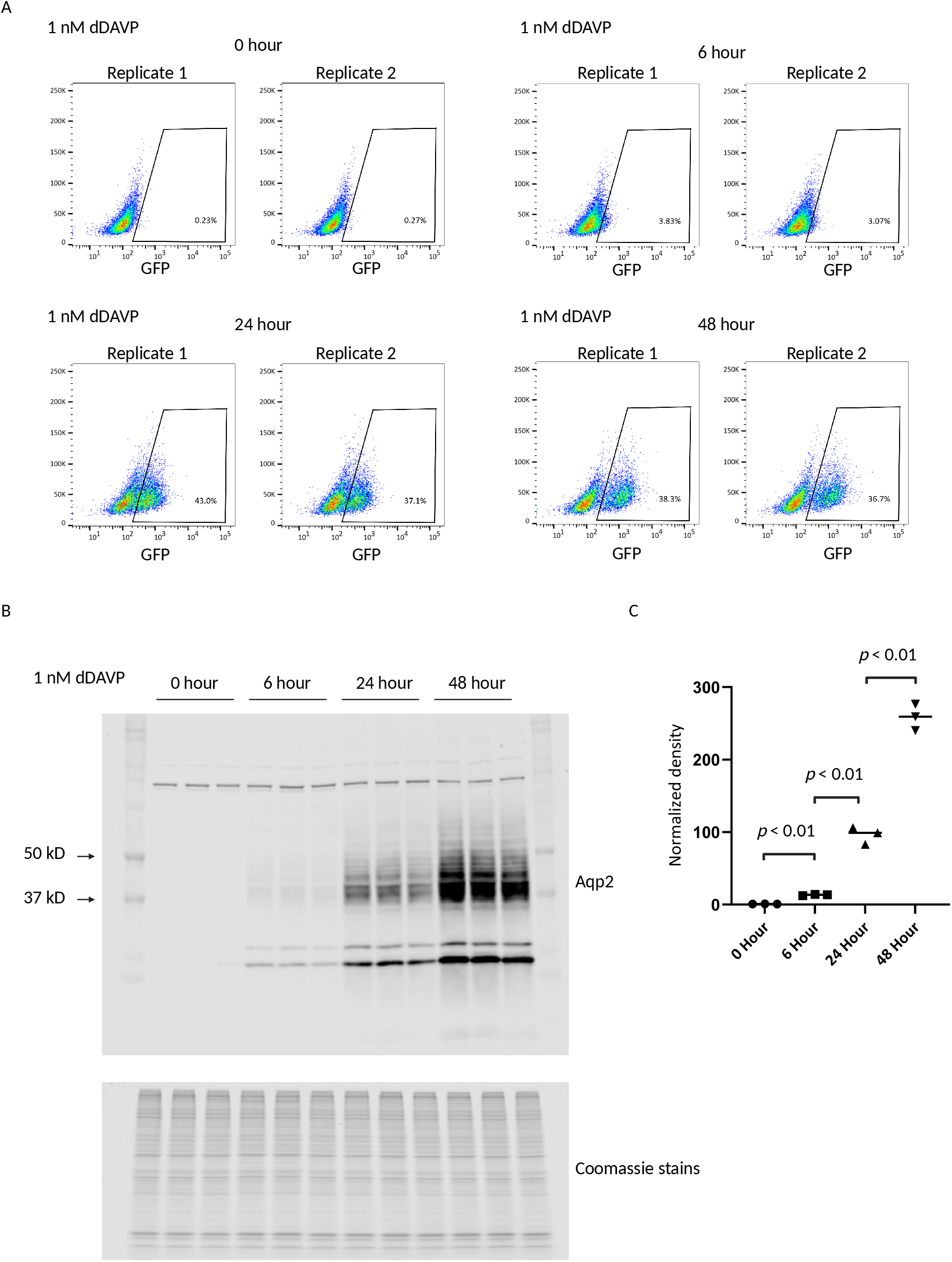
Time-dependent regulation of Aqp2 expression by dDAVP. *A:* FACS analysis of percentage of GFP-positive cells over 0, 6, 24, and 48 hours of 1 nM dDAVP treatment. Two replicates are present in each condition. *B:* Western blot comparing Aqp2 expression levels at the same four time points of dDAVP treatment. Coomassie stains serves as a loading control *C:* Quantitative analepsis of the Aqp2 expression from *B*. There is a significant time-dependent increase in Aqp2 expression (p < 0.01).

Aqp2 regulation is mediated by vasopressin binding to the V2 receptor (V2R), which activates adenylyl cyclase 6 (AC6) to increase intracellular cyclic-AMP (cAMP). This, in turn, activates protein kinase A (PKA), resulting in an upregulation of Aqp2 expression (33). Forskolin (FSK) also stimulates Aqp2 expression by directly activating AC6, but independent of V2R (34). To examine the effect of FSK on GFP expression, we performed FACS analysis on cells treated with either 1nM dDAVP or 10 uM FSK for 48 hours (untreated cells served as the control). As shown in Fig. 4, A and B, the proportion of GFP-positive cells increased significantly to 45.3% ± 0.98% in response to dDAVP and 50.8% ± 1.2% in response to FSK, compared to 1.04% ± 0.44% in the control group. The large proportion of cells expressing no GFP after stimulation further confirmed the significant variability. Both dDAVP and FSK increased the GFP intensity per cell, as indicated by the x-axis of the flow cytometry plots. These data further confirm the previous observations that our new GFP reporter system is a reliable tool for studying Aqp2 regulation and holds great potential for use in the high-throughput screening of Aqp2 regulators.

**Figure 4.**
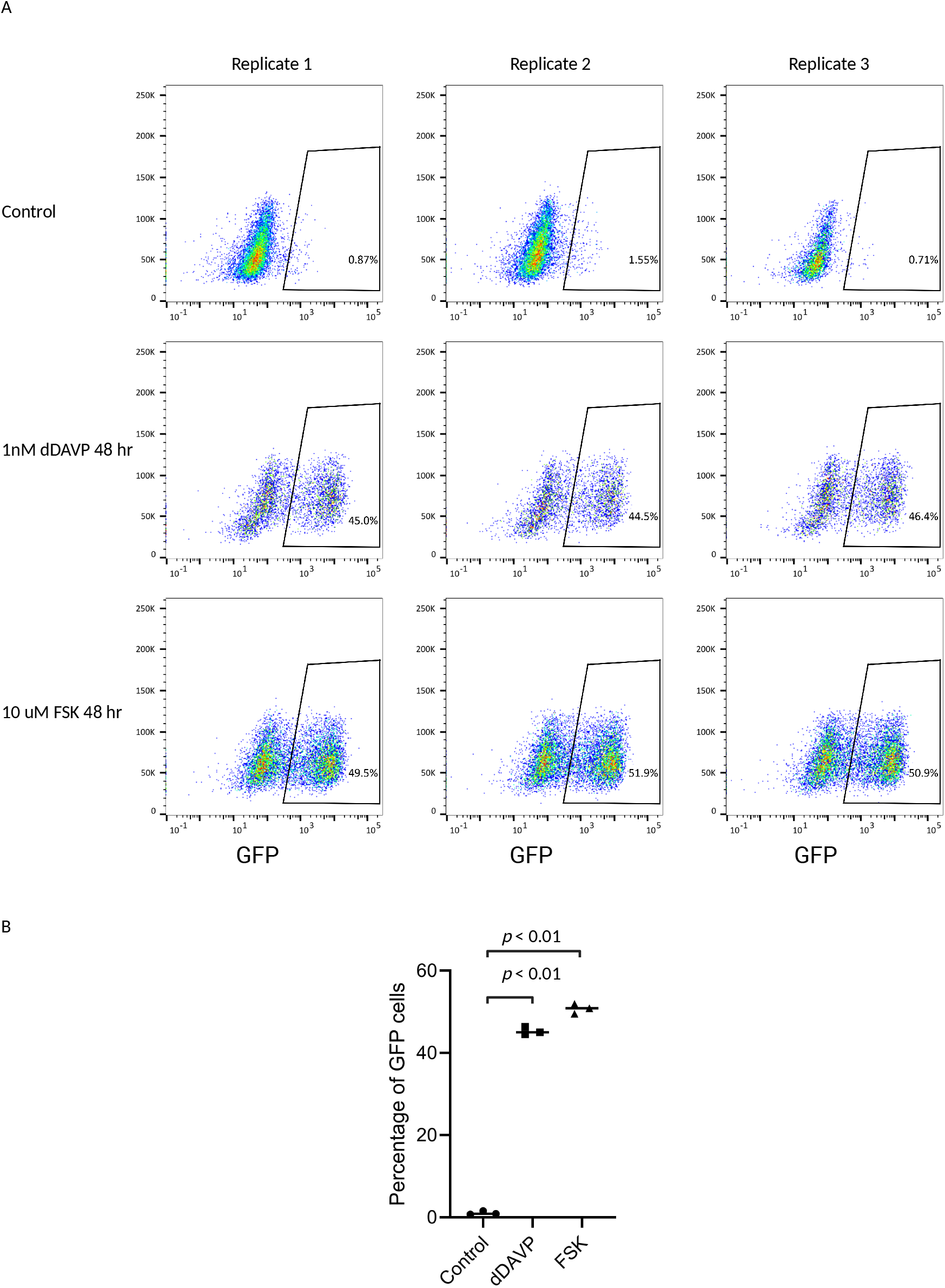
Both dDAVP and Forskolin result in increase in Aqp2-GFP expression. *A:* FACS analysis of dDAVP and Forskolin (FSK) response. FACS plots across three replicates to compare the effects of dDAVP (1 nM) and FSK (10 µM) on the GFP reporter expression after 48 hours of treatment. Cells without treatment server as control. *B:* Quantitative comparison of the percentage of GFP-positive cells across the three conditions. Similar to dDAVP, FSK significantly increases GFP-positive cells (p < 0.01).

### Heterogeneity in Aqp2 expression

To explore the basis of the observed heterogeneity, we FACS sorted the GFP^−^ cells (Aqp2^−^) and GFP^+^ (Aqp2^+^) cells, repopulated each group, and subsequently restimulated them separately with dDAVP (Fig. 5A). Interestingly, cells originating from GFP^−^ cells give rise to both GFP^−^ cells and GFP^+^ cells, and GFP^+^ cells similarly regenerate both GFP^−^ and GFP^+^ populations, yielding similar proportions of GFP^+^ and GFP^−^ cells. This result suggests that the two subpopulations are not terminally fixed but can dynamically interconvert, indicating an intrinsic plasticity in Aqp2 expression states.

**Figure 5.**
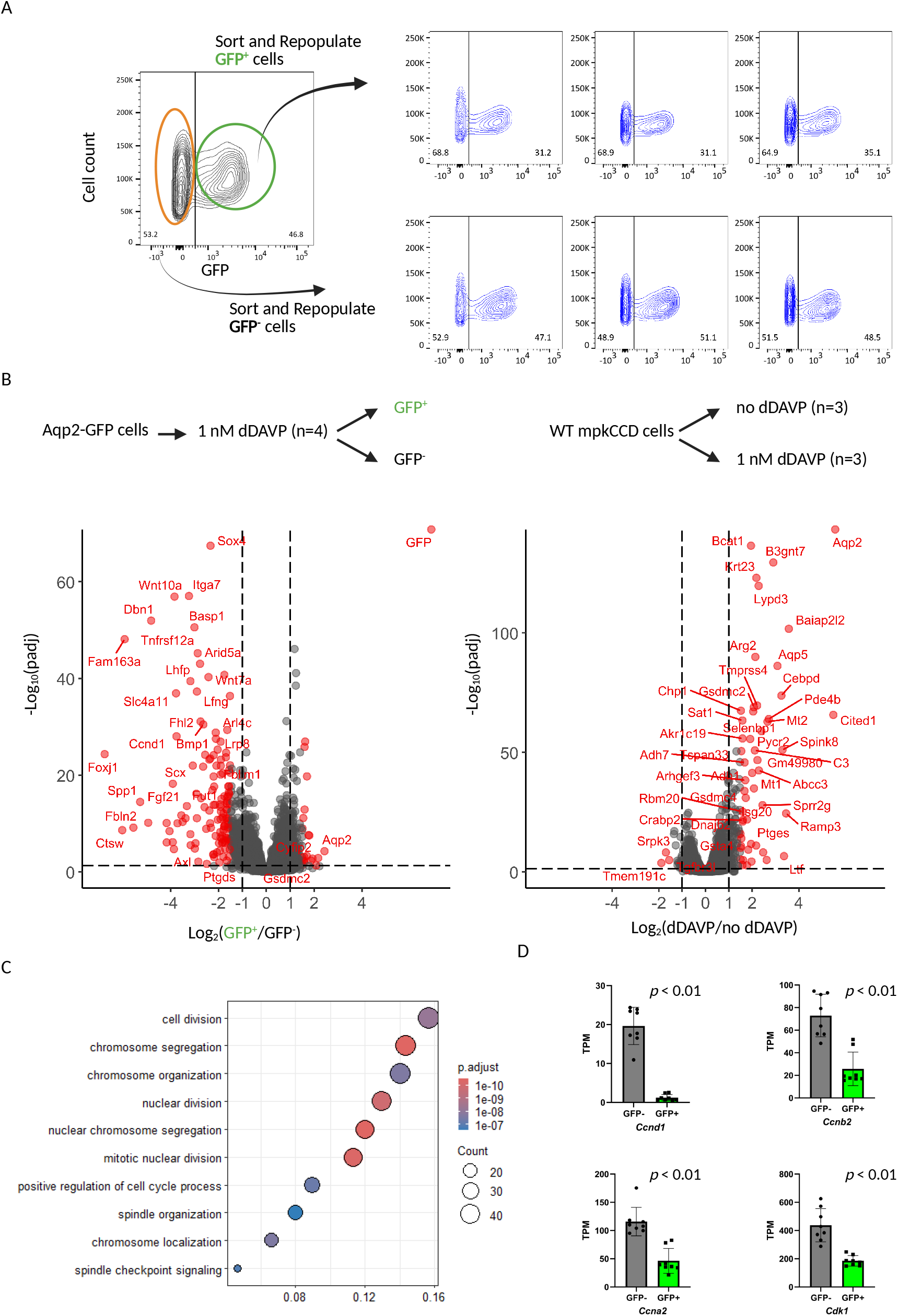
Heterogeneity in Aqp2-GFP expression and transcriptomic differences between GFP^+^ and GFP^−^ Aqp2 reporter cells. *A:* Flow cytometric analysis and repopulation experiments. Aqp2-GFP reporter cells were sorted into GFP^+^ and GFP^−^ subpopulations, which were then regrown and re-analyzed after dDAVP stimulation. Both sorted GFP^+^ and GFP^−^ cells regenerated heterogeneous populations, yielding similar proportions of GFP^+^ and GFP^−^ cells, indicating dynamic interconversion between the two states. Representative FACS plots show GFP distribution before and after sorting. *B:* Transcriptomic profiling of GFP^+^ versus GFP^−^ cells and dDAVP-stimulated versus untreated wild-type mpkCCD cells. Volcano plots highlight significantly differentially expressed genes (adjusted p < 0.01). GFP^−^ cells were enriched in transcripts associated with cell-cycle regulation (e.g., *Ccnd1, Ccnb2, Ccna2*, and *Cdk1*), whereas GFP^+^ cells exhibited higher Aqp2 expression. Parallel analysis of wild-type mpkCCD cells confirmed dDAVP-dependent induction of Aqp2 and other canonical target genes. *C:* Gene ontology (GO) enrichment analysis of transcripts upregulated in GFP^−^ cells. The most significant pathways were linked to cell division, chromosome segregation, nuclear division, spindle organization, and checkpoint signaling, consistent with the cell cycle association of GFP^−^ cells. *D:* Validation of selected cell cycle-related genes by RNA-seq quantification. Expression of *Ccnd1, Ccnb2, Ccna2*, and *Cdk1* was significantly elevated in GFP^−^ cells relative to GFP^+^ cells (all p < 0.01).

To investigate the molecular mechanisms of the underlying heterogeneity, we carried out RNA-seq on the sorted GFP^−^ cells (Aqp2^−^) and GFP^+^ (Aqp2^+^) cells, alongside the wild-type mpkCCD cells with or without dDAVP stimulation (Fig. 5B). Differentially expressed genes are shown by volcano plots and can be accessed at https://esbl.nhlbi.nih.gov/Reporter/. The full curated dataset is provided and downloadable on the same user-friendly web page. Not surprisingly, *GFP* and *Aqp2* are the two most enriched transcripts in the GFP^+^ cells compared to GFP^−^ cells, emphasizing the fidelity of the GFP in tracking Aqp2 transcriptional regulation. The analysis identified a number of transcripts that were highly enriched in the GFP^−^ cells. These includes *Wnt10a, Wnt7a*, and *Sox4*, which play crucial roles in development and are involved in the Wnt/β-catenin signaling pathway, as well as *Ccnd1* (Cyclin D1) which is a key regulator of the cell cycle. Consistent with prior findings, dDAVP stimulation in wild-type mpkCCD cells significantly upregulated *Aqp2* and other canonical vasopressin-responsive transcripts. The analysis did not find a significant difference in the abundance of the key markers (*Slc4a1, Slc26a4, Foxi1, Atp6v1c2, Atp6v1g3, Hmx2*, and *Dmrt2*) (35) associated with intercalated cells when comparing the GFP^−^ cells (Aqp2^−^) and GFP^+^ (Aqp2^+^) cells, suggesting there is no evidence of conversion of principal-like cells to intercalated-like cells. To systematically characterize the differences between GFP^−^ and GFP^+^ cells, we performed gene enrichment analysis. Upregulated transcripts in GFP^−^ cells were strongly associated with cell division, chromosome segregation, and checkpoint signaling (Fig. 5C). Consistently, expression of some of the cell-cycle related transcripts were significantly elevated in GFP^−^ cells (*Ccnd1, Ccnb2, Ccna2*, and *Cdk1*) compared to GFP cells (Fig. 5D).

To directly investigate the relationship between cell cycle status and Aqp2 expression, we performed cell cycle analysis using FACS (4⍰,6-diamidino-2-phenylindole [DAPI] staining). Indeed, we found GFP^+^ show less DNA content (DAPI intensity) compared to GFP^−^ cells (Fig. 6). Together, these findings indicate that Aqp2 heterogeneity in mpkCCD cells is at least partly driven by differences in cell cycle phase, with proliferative GFP^−^ cells displaying reduced Aqp2 expression, whereas quiescent GFP^+^ cells exhibit higher Aqp2 levels.

**Figure 6.**
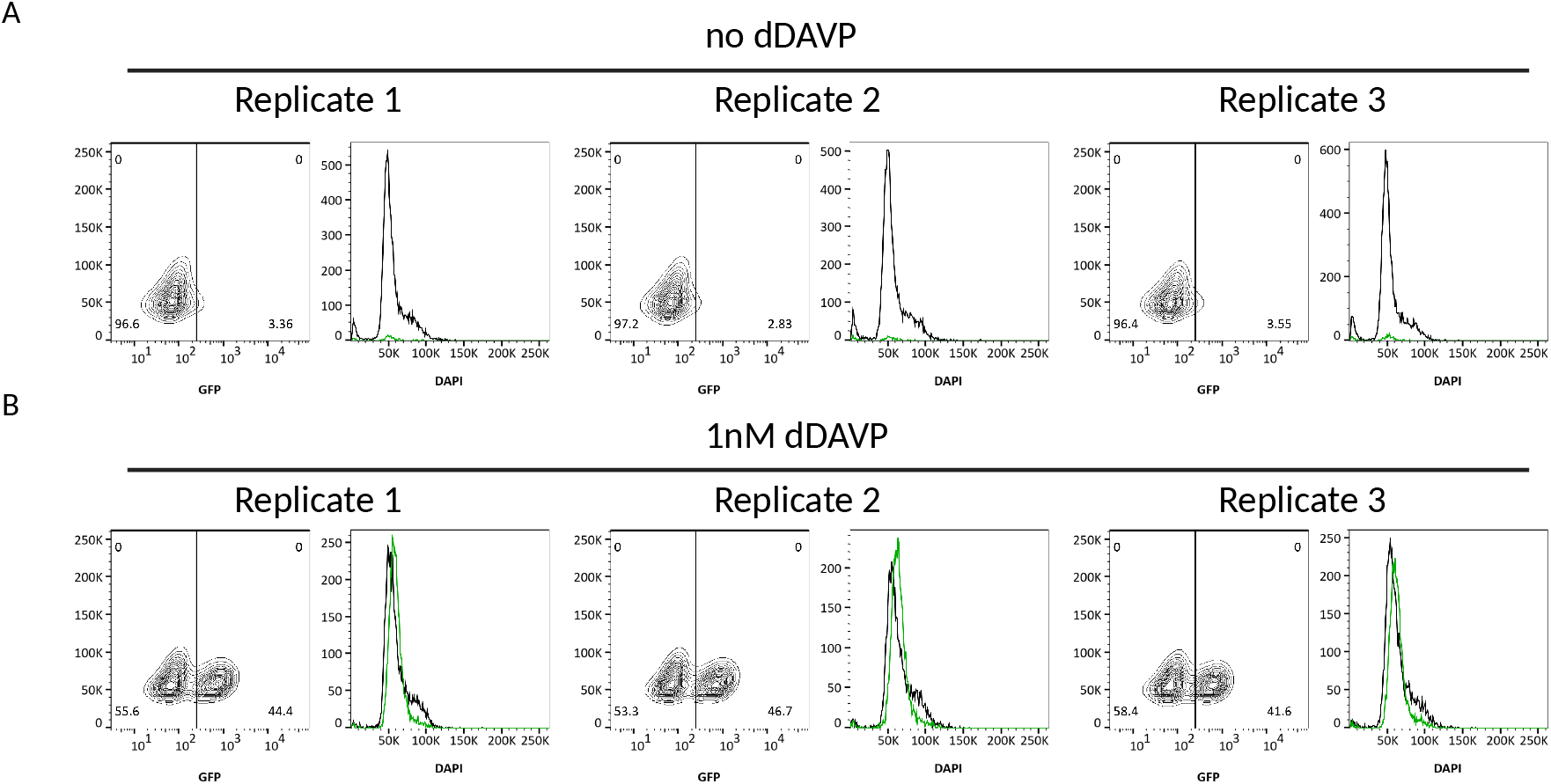
Cell cycle differences between dDAVP-responsive and non-responsive cells. *A:* Without dDAVP treatment, the Aqp2-GFP reporter cells have minimum GFP expression and most of the cells with the DAPI staining peaked at the intensity around 50K (G0/G1). Some cells are in the G2/S phase with intensity around 50K∼100K. *B:* With dDAVP stimulation, the cells with GFP expression (dDAVP responsive cells, green curve) show less DNA content at the density around 100K; the cells without GFP expression (dDAVP non-responsive cells, black line) show similar DNA content profile in *A*.

## DISCUSSION

In this study, we successfully engineered an endogenously driven Aqp2-GFP reporter in mpkCCD cells using CRISPR/Cas9 technology. The GFP was inserted at the endogenous *Aqp2* locus, ensuring the reporter is regulated by the native *Aqp2* promoter. Extensive characterization—including immunoblotting, immunofluorescence, FACS analysis, and kinetic studies—confirmed that the resulting cell line faithfully and dynamically tracks *Aqp2* transcriptional regulation by vasopressin (dDAVP) and forskolin (FSK). The simultaneous and concordant increases in both Aqp2 and GFP mRNA and protein expression validate this system as a reliable and valuable tool for studying the mechanisms governing Aqp2 expression. The full curated dataset is provided and downloadable on a user-friendly web page at https://esbl.nhlbi.nih.gov/Reporter/.

The design of this reporter system provides several key advantages. By targeting the endogenous locus, we maintain the native regulatory structure, which is often lost in traditional exogenous (plasmid) systems. Furthermore, we provide a detailed experimental scheme that is broadly applicable to generating other endogenous reporter systems in diverse cell line and gene models. Most significantly, this approach and the validated components can be readily adapted for *in vivo* use to create a native *Aqp2* reporter mouse model. Such a model, which faithfully mimics the endogenous expression pattern of Aqp2, is currently lacking in the field, especially in developmental biology.

In the present study, the insertion of the P2A peptide separates GFP from Aqp2, meaning this cell line is not suitable for use of the GFP signal in microscopic studies of Aqp2 trafficking to the apical membrane, as the GFP is restricted to the N-terminus and is cleaved off post transcriptionally. While we observed a timely concordant expression of the reporter and native protein, the differences in protein half-lives (e.g., GFP at 26 hours vs. Aqp2 at 14 hours following dDAVP stimulation) (36) suggest that the GFP signal may not fully capture rapid or transient changes in Aqp2 protein levels during short-term stimuli or removal of the stimulus. Researchers must consider these differences when interpreting kinetic data. Although dDAVP is typically used at 0.1 nM to mimic physiological conditions, we increased the concentration to 1 nM in the present study to achieve a more robust transcriptional response.

The reporter cell line provides an opportunity to investigate the cell-to-cell heterogeneity in Aqp2 expression. Our FACS analysis demonstrated that even after maximal dDAVP or FSK stimulation, less than half of the cells express Aqp2/GFP, confirming a high degree of intrinsic variability. Cell-sorting experiments revealed that the GFP^−^ (Aqp2^−^) and GFP^+^ (Aqp2^+^) populations are not terminally fixed. Upon re-stimulation, both sorted populations were able to regenerate the heterogeneous mix, indicating a remarkable intrinsic plasticity or dynamic interconversion between the Aqp2-expressing and non-expressing states. This observation aligns with reports of cellular heterogeneity and plasticity in other biological systems (37) and native kidney cells (38). The mechanisms underlying this variability remain poorly understood, but several possible explanations have been proposed. One possibility is that acquired genetic mutations during the prolonged cell culture may prevent Aqp2 expression. This is likely not the case, as the phenotype is reversable. Another hypothesis is that a subset of mpkCCD cells may undergo phenotypic transition from the principal cell-like state to intercalated cell-like cells, as seen in native kidney collecting duct (39). However, examination of the specific markers associated with intercalated cells (*Slc26a4, Slc4a1, Foxi1*, and others) did not reveal significant differences when comparing the GFP-negative and -positive cells. Finally, incomplete differentiation and differences in cell cycle phase can also contribute to the lack of Aqp2 expression in response to dDAVP. Studies have shown that cell cycle regulation may contribute to the cellular variation (40). Indeed, RNA-seq analysis of the GFP^−^ (Aqp2^−^) and GFP^+^ (Aqp^+^2) cells revealed significant enrichment of cell cycle-related transcripts in the GFP^−^ cells, including cyclins and cyclin-dependent kinases such as *Cdk1, Ccnd1, Ccnb2*, and *Ccna2*. Gene Ontology (GO) analysis confirmed this finding, with terms like “cell division” and “chromosome segregation” being highly represented in the GFP^−^ population. Direct FACS analysis of DNA content further supported these findings, demonstrating that the GFP (Aqp2-expressing) cells are predominantly in the G0/G1 phase (quiescent), while a greater proportion of the GFP^+^ (non-expressing) cells are actively engaged in the G2/S phases (proliferative). This strongly suggests that Aqp2 expression is inversely coupled to the cell cycle—a finding consistent with previous observations in kidney collecting duct cells that link increased proliferative programs to decreased Aqp2 expression (41). In addition to cell cycle difference, several transcription factors (including Sox4, Gata3, and others) and ligands (*Wnt10a* and *Wnt7a*) for Wnt/β-catenin pathway show difference between these two populations. Their direct roles in Aqp2 expression and cell plasticity remain further investigation.

One of the emerging methodologies in identifying novel genes involved in complex biological systems is phenotypic screens. Our reporter line paves the way for future high-throughput screening for regulators of *Aqp2* gene transcription. The ability to quantify Aqp2 expression via GFP fluorescence in a scalable format makes this system ideal for: 1) drug screening to identify small molecules that can either enhance Aqp2 expression (e.g., for certain forms of Nephrogenic Diabetes Insipidus, NDI) or suppress it; 2) CRISPR-mediated functional screens (e.g., using CRISPRa/i or knockouts) targeting specific gene families, such as kinases or transcription factors, to systematically map both positive and negative regulators of Aqp2 transcription. Ultimately, the combination of this robust reporter system with new techniques will be pivotal in dissecting the complex regulatory network underlying Aqp2 expression and the maintenance of water balance.

## DATA AVAILABILITY

The RNA-seq results are accessible on a publicly accessible webpage at https://esbl.nhlbi.nih.gov/Reporter/. RNA seq raw data are deposited in https://www.ncbi.nlm.nih.gov/geo/query/acc.cgi?acc=GSE310089.

## SUPPLEMENTAL MATERIAL

This article contains supplemental material at https://doi.org/10.25444/nhlbi.30794933.

**Supplemental Figure 1**. Gene structure and DNA sequence of the Aqp2-reporter.

**Supplemental Figure 2**. Additional western blotting of Aqp2-reporter clones.

**Supplemental Table 1**. RNA-seq analysis of the dDAVP response of mpkCCD cells and GFP-negative cells and GFP-positive cells.

## ACKNOWLEDGEMENTS

We utilized the NHLBI Flow Cytometry Core Facility (Pradeep Dagur, Director), The NHLBI DNA Sequence Core (Yuesheng Li, Director) and the NHLBI Light Microscopy Core Facility (Christian Combs, Director) for these studies.

## GRANTS

The work was funded by the Division of Intramural Research, National Heart, Lung and Blood Institute [project ZIA-HL001285 and ZIA-HL006129, M.A.K.]. The content is solely the responsibility of the authors and does not necessarily represent the official views of the National Institutes of Health.

## DISCLOSURES

The authors declare that they have no conflicts of interest with the contents of this article.

## AUTHOR CONTRIBUTIONS

Lihe Chen and Mark A. Knepper conceived and designed research, Lihe Chen performed experiments, Lihe Chen and Mark A. Knepper analyzed data, Lihe Chen, Adrian Rafael Murillo-de-Ozores, Euijung Park, Shuo-Ming Ou, and Mark A. Knepper interpreted results of experiments, Lihe Chen prepared figures, Lihe Chen and Mark A. Knepper drafted manuscript, Lihe Chen and Mark A. Knepper edited and revised manuscript, Lihe Chen, Adrian Rafael Murillo-de-Ozores, Euijung Park, Shuo-Ming Ou, and Mark A. Knepper approved final version of manuscript

## Notes

https://doi.org/10.25444/nhlbi.30794933

https://esbl.nhlbi.nih.gov/Reporter/

### Competing Interest Statement

The authors have declared no competing interest.

## REFERENCES

1. Snijder B, and Pelkmans L. Origins of regulated cell-to-cell variability. Nat Rev Mol Cell Biol 12: 119–125, 2011.

2. West-Eberhard MJ. Developmental plasticity and the origin of species differences. Proc Natl Acad Sci U S A 102 Suppl 1: 6543–6549, 2005.

3. Moczek AP, Sultan S, Foster S, Ledon-Rettig C, Dworkin I, Nijhout HF, Abouheif E, and Pfennig DW. The role of developmental plasticity in evolutionary innovation. Proc Biol Sci 278: 2705–2713, 2011.

4. Marusyk A, and Polyak K. Tumor heterogeneity: causes and consequences. Biochim Biophys Acta 1805: 105–117, 2010.

5. Dagogo-Jack I, and Shaw AT. Tumour heterogeneity and resistance to cancer therapies. Nat Rev Clin Oncol 15: 81–94, 2018.

6. Longley DB, and Johnston PG. Molecular mechanisms of drug resistance. J Pathol 205: 275–292, 2005.

7. Gawad C, Koh W, and Quake SR. Single-cell genome sequencing: current state of the science. Nat Rev Genet 17: 175–188, 2016.

8. Muzzey D, and van Oudenaarden A. Quantitative time-lapse fluorescence microscopy in single cells. Annu Rev Cell Dev Biol 25: 301–327, 2009.

9. Pearce D, Soundararajan R, Trimpert C, Kashlan OB, Deen PM, and Kohan DE. Collecting duct principal cell transport processes and their regulation. Clin J Am Soc Nephrol 10: 135–146, 2015.

10. Knepper MA, Kwon TH, and Nielsen S. Molecular physiology of water balance. N Engl J Med 372: 1349–1358, 2015.

11. D’Acierno M, Fenton RA, and Hoorn EJ. The biology of water homeostasis. Nephrol Dial Transplant 40: 632–640, 2025.

12. Olesen ETB, and Fenton RA. Aquaporin 2 regulation: implications for water balance and polycystic kidney diseases. Nat Rev Nephrol 17: 765–781, 2021.

13. Cheung PW, Bouley R, and Brown D. Targeting the Trafficking of Kidney Water Channels for Therapeutic Benefit. Annu Rev Pharmacol Toxicol 60: 175–194, 2020.

14. Kortenoeven ML, Pedersen NB, Rosenbaek LL, and Fenton RA. Vasopressin regulation of sodium transport in the distal nephron and collecting duct. Am J Physiol Renal Physiol 309: F280–299, 2015.

15. Ecelbarger CA, Kim GH, Terris J, Masilamani S, Mitchell C, Reyes I, Verbalis JG, and Knepper MA. Vasopressin-mediated regulation of epithelial sodium channel abundance in rat kidney. Am J Physiol Renal Physiol 279: F46–53, 2000.

16. Bockenhauer D, and Bichet DG. Pathophysiology, diagnosis and management of nephrogenic diabetes insipidus. Nat Rev Nephrol 11: 576–588, 2015.

17. Nielsen S, Frokiaer J, Marples D, Kwon TH, Agre P, and Knepper MA. Aquaporins in the kidney: from molecules to medicine. Physiol Rev 82: 205–244, 2002.

18. Valenti G, and Tamma G. The vasopressin-aquaporin-2 pathway syndromes. Handb Clin Neurol 181: 249–259, 2021.

19. Verbalis JG. Disorders of water metabolism: diabetes insipidus and the syndrome of inappropriate antidiuretic hormone secretion. Handb Clin Neurol 124: 37– 52, 2014.

20. Sandoval PC, Claxton JS, Lee JW, Saeed F, Hoffert JD, and Knepper MA. Systems-level analysis reveals selective regulation of Aqp2 gene expression by vasopressin. Sci Rep 6: 34863, 2016.

21. Hasler U, Mordasini D, Bens M, Bianchi M, Cluzeaud F, Rousselot M, Vandewalle A, Feraille E, and Martin PY. Long term regulation of aquaporin-2 expression in vasopressin-responsive renal collecting duct principal cells. J Biol Chem 277: 10379–10386, 2002.

22. Yu MJ, Miller RL, Uawithya P, Rinschen MM, Khositseth S, Braucht DW, Chou CL, Pisitkun T, Nelson RD, and Knepper MA. Systems-level analysis of cell-specific AQP2 gene expression in renal collecting duct. Proc Natl Acad Sci U S A 106: 2441–2446, 2009.

23. Hsu PD, Lander ES, and Zhang F. Development and applications of CRISPR-Cas9 for genome engineering. Cell 157: 1262–1278, 2014.

24. Dobin A, Davis CA, Schlesinger F, Drenkow J, Zaleski C, Jha S, Batut P, Chaisson M, and Gingeras TR. STAR: ultrafast universal RNA-seq aligner. Bioinformatics 29: 15–21, 2013.

25. Li B, and Dewey CN. RSEM: accurate transcript quantification from RNA-Seq data with or without a reference genome. BMC Bioinformatics 12: 323, 2011.

26. Love MI, Huber W, and Anders S. Moderated estimation of fold change and dispersion for RNA-seq data with DESeq2. Genome Biol 15: 550, 2014.

27. Bens M, Vallet V, Cluzeaud F, Pascual-Letallec L, Kahn A, Rafestin-Oblin ME, Rossier BC, and Vandewalle A. Corticosteroid-dependent sodium transport in a novel immortalized mouse collecting duct principal cell line. J Am Soc Nephrol 10: 923– 934, 1999.

28. Dambournet D, Hong SH, Grassart A, and Drubin DG. Tagging endogenous loci for live-cell fluorescence imaging and molecule counting using ZFNs, TALENs, and Cas9. Methods Enzymol 546: 139–160, 2014.

29. Paquet D, Kwart D, Chen A, Sproul A, Jacob S, Teo S, Olsen KM, Gregg A, Noggle S, and Tessier-Lavigne M. Efficient introduction of specific homozygous and heterozygous mutations using CRISPR/Cas9. Nature 533: 125–129, 2016.

30. Liu Z, Chen O, Wall JBJ, Zheng M, Zhou Y, Wang L, Vaseghi HR, Qian L, and Liu J. Systematic comparison of 2A peptides for cloning multi-genes in a polycistronic vector. Sci Rep 7: 2193, 2017.

31. Lin S, Staahl BT, Alla RK, and Doudna JA. Enhanced homology-directed human genome engineering by controlled timing of CRISPR/Cas9 delivery. Elife 3: e04766, 2014.

32. Koch B, Nijmeijer B, Kueblbeck M, Cai Y, Walther N, and Ellenberg J. Generation and validation of homozygous fluorescent knock-in cells using CRISPR-Cas9 genome editing. Nat Protoc 13: 1465–1487, 2018.

33. Chen L, Jung HJ, Datta A, Park E, Poll BG, Kikuchi H, Leo KT, Mehta Y, Lewis S, Khundmiri SJ, Khan S, Chou CL, Raghuram V, Yang CR, and Knepper MA. Systems Biology of the Vasopressin V2 Receptor: New Tools for Discovery of Molecular Actions of a GPCR. Annu Rev Pharmacol Toxicol 62: 595–616, 2022.

34. Maric K, Oksche A, and Rosenthal W. Aquaporin-2 expression in primary cultured rat inner medullary collecting duct cells. Am J Physiol 275: F796–801, 1998.

35. Chen L, Lee JW, Chou CL, Nair AV, Battistone MA, Paunescu TG, Merkulova M, Breton S, Verlander JW, Wall SM, Brown D, Burg MB, and Knepper MA. Transcriptomes of major renal collecting duct cell types in mouse identified by single-cell RNA-seq. Proc Natl Acad Sci U S A 114: E9989–E9998, 2017.

36. Sandoval PC, Slentz DH, Pisitkun T, Saeed F, Hoffert JD, and Knepper MA. Proteome-wide measurement of protein half-lives and translation rates in vasopressin-sensitive collecting duct cells. J Am Soc Nephrol 24: 1793–1805, 2013.

37. Meacham CE, and Morrison SJ. Tumour heterogeneity and cancer cell plasticity. Nature 501: 328–337, 2013.

38. Chang-Panesso M, and Humphreys BD. Cellular plasticity in kidney injury and repair. Nat Rev Nephrol 13: 39–46, 2017.

39. Wu H, Chen L, Zhou Q, Zhang X, Berger S, Bi J, Lewis DE, Xia Y, and Zhang W. Aqp2-expressing cells give rise to renal intercalated cells. J Am Soc Nephrol 24: 243–252, 2013.

40. Buenrostro JD, Wu B, Litzenburger UM, Ruff D, Gonzales ML, Snyder MP, Chang HY, and Greenleaf WJ. Single-cell chromatin accessibility reveals principles of regulatory variation. Nature 523: 486–490, 2015.

41. de Groot T, Alsady M, Jaklofsky M, Otte-Holler I, Baumgarten R, Giles RH, and Deen PM. Lithium causes G2 arrest of renal principal cells. J Am Soc Nephrol 25: 501–510, 2014.

