## Supplemental Figure 1 for "GFP Reporter System Reveals Cell-to-Cell Variability in Aquaporin-2 Expression"

A

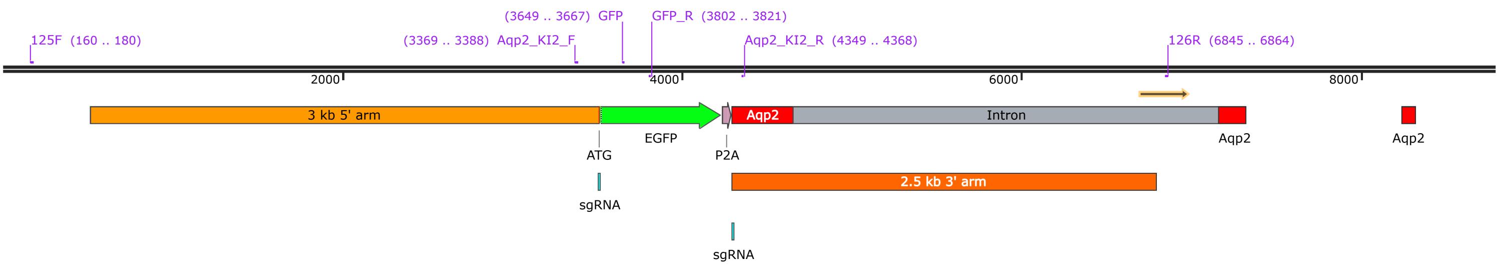


B

TGCCTACAAAGTGCATGAGCTGTGTCCAGAACCAAGGCTGAGCAGTCAAGTCTCACAATGGAGCACAGCTGACCCCACCAAGAACGCTTGGAATCCATCAGACGTTTGATTTTGATTGTTGTTGTGCATTCAG
AAACCTTTTACCGTGGCCAAGGTTGGCGTGAGCCCCTCATTCAGTTCTGTGACCCACAATTTGCGAGGCTGGAAGGGTAAAGATAAACAAAAGCAAAAGCAAACACGGGAGGATACGGCGGCGGCCTCC
GGGAGCATGCGAACCCACAGCCGCAGCTGGGCCTCCGGTAGGGATGCTGCTGCTGCTGCTTCCCTGGCATGAAGGAAGGCTGAGTACAAAGCCGGGGCATAGACCATGCGGTTATGGCTGAACTCCCC
CTCCCCGCTGGCTTGGGATCAGACCAGAATATTCTCACCGCTCCTGGAAGTCAGTCTCCGGGCATCATGTTCTAGTCTGTGAAATGAGACCCATGGCTATCATGGTCTGGCTGATCCCATCCTTGCTTTTTTGT
TTTACTTGTTGTTGTTGTTGTTGTTGTTGTTGTTGTTGTTGTTGCTGCTGCTGCTGCTGCTGCTGCTGCTGCTGCTGCTGCTGCTGCTGCTGCTGCTGCTGCTGCTGTTTTCGGCACTGGGAATGGAACCCAGGAT
TTACTACCATGTCCTGGGCAAGTGCTCTTCCACAGAACCACACCTCCCGCACCGTGGAAATGTGGATGTATGGCAAAACCACACAAATTGGTGTGTGGTTTCCTGGGCTGTACAGAAGCAGCGACACCTATT
ATGGGTCTACCATGACGCCAGGTTGGTTAGAGGCGGTGCTGTTTAGGGTCAGAACCAGTGACTGCTGTTCTCATGATCATACACAGTGTATAGATCAGAAAGGGTGTGTGGCTCACCAAAGGTCACTCAGGAA
CTCGATAGTCAACGGTGGGTTTGGAACCACTGTCACAGTTCAGCTCAATATGGCCTGGGGGTTTTCCAGGTGCCCCTGAGCCAGGCAGGGACTGGGCACATGGAGGGTGGAAGATACTACATACAAGATAG
TATGTATCATCAAGATAAAACATGATACATTCCCTGGGTATGGTAGACAGTACATACAAGATAGTATGTATCATCAAGGCCCAATCTTGATACCCTCCACCCTTTCTGGGTGTGCTATGGTTGAAGAAGAAAACAGA
GTCACAGGTACCCTTCTCAAGGCAAGATTCTTCCACCCTCCCTCTGAGTCTTTCTGGAGAGCGTGGATTGCAGGGGTGGGGGATGGAAGTGGGGAGGAGGACCCTGAACAGGGAGTACTACCTGAGCCCA
CATCCCCAGAGGCTCCTCTGAGCTGGCCTCTGTTGGGGTCATGTATTGTTTCTTCCTCCTCATTCTTGTCTTTCCCCCTAATGGCAAGAGTTGCTAGTCATGTGGGGGTCGGGGTAAATGTGCATGGACCCGTG
TTGTGTATGTGGGGGATTTTTGTGCATCTTACTACATTCTGAAAACAAATGTATGGGAGGGACATCACTGCCATTGCTGTCTGGGCGAGGAGACCAGGGCTCAGAGCATGCCTGGGGAGTGAGGGTGTCAGGA
AAGCACTCCTCTCTAGGTTCTAGCCTGAGCTCAGCCACCACTCTTCTGGGGCACCTTGGGTGAGTAACAGGCCCTCCCTTGGCTTCAGTCACCCCTTAGTGTAAGTATCTCTATCCTACGATCATCGGTTTCTC
CCACATTCCTGTTCCTTGGATTGTGGCTGGCCAGGAGGACTAGAAGCCCGCCATCTGGATGCACTGTAGGCCCTCCTGCCTTTGGATCCCCAAAGAAAGAAGAGGCGTAGGGTAGGTCCGTCTTCTGAGC
CTCAACAGCCATCAGAGCTGTGACAGCAGTGGCCTTGCAGAACACTGACCAAGCTTAGTGGCACCAAGACTAGAGGTTACTAGCAGGCACCATGGCCTCCTTGAGCCTGGAGAGCTAGCTGCAACTCCAG
AAACATCTTCCTCCTGTGGTAAGGGTGGCTCTGAGCTCGAGATGGTATCTAGAGCCTGTTGTAGCCTTGGAGACTCTGAGTTTGGGGGAGACCTGTTTGCCTCAAAATGCAAATAGCACTGAGCAATAGCTGA
CAAAACCTGAGTTCTTCCTGAGAACCAGGAGCCATATTAATGCAAATCTCTGTCCTATGAGATAGTTTTATTATCTCATGTACACAGGCAGAGCAGAACTGTCACATCTCAACGACATGAGTGTGTGCTGCAGAA
CAGAAACCCTCATGTTCCTAAGTTGACCTTGGTGACTCTATCACGCAGGGCTAGGGTCCCAGGGCCACTGAAGCCACCTTTGGCTCCAGGCAATGGGAGTTCCTCTCTGGCTGAGATTCCTGGGTGAAGGT
GGAACATAATGACTTATGGGTGATTAACTGCAAGAAAGCAAGCCCAAGACCTTTTGCCTTGGAAATTTGTCCCAGGCCAGCCAGTTGTTTTCCATCCTCTTCCAGAAGAAAGACCATCCAGTCCGCGGAAAG
GCAGCGCCCACATTTCCTCACAGTTTGGGGTAAGGCATTGCCCCTTCCTTGGCCAGCCTTTTAGTCAAAGAGAAGGTCACTGGACACAGCCTCCTCTGCAGGAACTGGTGCTGGTGGTCATTGTGGGGGCT
GGGGCAGCCCTGAGGCAGCTCCATGGGGTAACTGAGGAAAAAACGAGGAAAACAGAGACGTCAATCCTTATCTGGAGTCCATTAATGAGGAGAACATTAGTCAGCTGTGAAAGCTAAGATGGGGTGATAGG
CCTTCGGGTGGGCAGGAGCAGGGATGGGGGCAAGTCCGCCATGGGGGAGCACAGGGTTGGCAGGAACGCCTCCTCCACCCCACGTGCCCAGGTCTACGATAGGAAGGCCCTATAAGTGCCCACAGT
CTAGCCTCTCCGGAGGCCCAGAGGAAGAGAGAAGAGAAAGAGAGAGGGAGGGAGGAAGAGCCACCCCCGTGGCCCAGACCACTGGCCAGCGCTCAGAAGGCCGAGCAGC**ATG**gtgagcaagggcgagg
agctgttcaccggggtggtgcccatcctggtcgagctggacggcgacgtaaacggccacaagttcagcgtgtccggcgagggcgagggcgatgccacctacggcaagctgaccctgaagttcatctgcaccaccggcaagctgcccgtgccctggcccaccctcgt
gaccaccctgacctacggcgtgcagtgcttcagccgctaccccgaccacatgaagcagcacgacttcttcaagtccgccatgcccgaaggctacgtccaggagcgcaccatcttcttcaaggacgacggcaactacaagacccgcgccgaggtgaagttcgagg
gcgacaccctggtgaaccgcatcgagctgaagggcatcgacttcaaggaggacggcaacatcctggggcacaagctggagtacaactacaacagccacaacgtctatatcatggccgacaagcagaagaacggcatcaaggtgaacttcaagatccgccacaa
catcgaggacggcagcgtgcagctcgccgaccactaccagcagaacacccccatcggcgacggccccgtgctgctgcccgacaaccactacctgagcacccagtccgccctgagcaaagaccccaacgagaagcgcgatcacatggtcctgctggagttcgt
gaccgccgccgggatcactctcggcatggacgagctgtacaagggaagcggagctactaacttcagcctgctgaagcaggctggagacgtggaggagaaccctggacctTGGGAACTCCGGTCCATAGCGTTCTCCCGAGCGGTGCTGGCCG
AGTTCCTGGCCACGCTCCTTTTCGTCTTCTTTGGCCTTGGCTCAGCCCTCCAGTGGGCCAGCTCCCCACCCTCTGTGCTCCAGATTGCCGTGGCCTTTGGTCTGGGCATTGGCACCCTGGTTCAGGCTCTG
GGCCATGTCAGCGGGGCCCACATCAACCCTGCTGTGACTGTGGCGTGCCTGGTGGGTTGCCATGTCTCCTTCCTTCGAGCTGCCTTCTACGTGGCTGCCCAGCTGCTGGGGGCCGTGGCCGGGGCCG
CCATCCTCCATGAGATTACCCCTGTAGAAATCCGCGGGGACCTGGCTGTCAATGCTGTGAGTAGCCCGAGTTTCATCCCTTCTGGGAGGGGCAGACACTGAAAACCCTTGTGAGGGGTGAGGTGGGTGGG
ACAGACTCTGAGCAGGGGGGTGGGGGAGTAAACAGAGAGCCGAAGGCCAGGAACACAGCTACCCAAGGAGAGCAACAGAGTCACCCGGAGCATCGAGTGTGGGGCCTTCCACCCAGCACCACCAAG
GAAAAGCAGCCTCGGTTTCTGTTAGACTCCATAGGAGGACTAGGCTAGACTGTAGCAGGACACAGCAACCTAGCACAGACAGGCCTTTGGGCCCTCGGGGACTCCAGGGAGCAGCGGCAAGGATCTAGA
GAGGAAGGGATGAGACATCCCAGCAGCACTGAGCATGGGAGGACTGGATAGGGCCCCTCACAAGGCAACTGCTGACAGCTCTGTGTCTGGGGGCAGGTTCCAAGCAGGGAGCGCCGCTAATCCCCCA
GACCCTGATTCAACCACAAAACCCATAATCTCTGCCTCAACCAACTATCTGAGGTGCCATATGGGCTTTGTCTCACCAACCAAGAAGCAGACTTGGAAAGGGAGACGGGAGATCAGATGTGCTCATGTGTTAA
TATCGGAAGTTCCAGAACCTAGCAGAAGCCTGCTGGAAGAGACTCTTGGAGTCTGCCTTCCTGTTATAATGGTGAACAGCACCTGGGGCACTTTGAGAGTTGAGGGTGAAGCCCTGGTCTCCACGCTCTGCA
GATCTAGAAATCTGGGGACCCCTTCAGCACAGCACGCACTCTGTACCCCAAACCAGGAAAGCAGGTGCCTCTCAGGCCTCCTCCTTGTCCTTCATTTCTCCCCTGCCTCAATCCCTTGGCCCTGCCCACA
CCCCCCTCTGCACGTGGCTTTGCTCATGAGAGGGACGGAGGCCCTCGGGTGAGTGAGCACCCTTTTCAAAAAATCAGAAACCATGGTGAGGCTGGACACCTAGAGCCAAAGAGGAACAAGGGGCCTGC
TGAGGGGTGGAGTGTTAGCCATGTCTATTAGAAAATAAATGAGCTTGTTTTACAACAGGGAGACACGCAGCTAGTCAAAGAACCCCCTGAAAACATACCGTAGGCTTTCTCTAGCAGTAAGGTGCTGGGAGCA
CTAAAGCCCTGGAGAATAGCCACCCCTGGTTCCCCCACCATATCCGAGCCTACCCAACCCTCAGCACCAAAGGCCGTTTCTCTGCATTGACCCACCTTAACTGGTAGAGGGGCCCCTGCCTGTCTGGAAT
AAGGCATCAGAGGGCCCATTTGCCCTGAGGCTAGTCAGAGACCAGTCAGGTGTTCTCTGAGGATGCTATCAAGAGTCAGTTTGGACCCCAGGAAGCTGCCACCAATGGATAGCGTCCCTGTGCCAAGCCA
CATACTATCCTATCCTTGGTAGACCACTGTAAGTTTAGTTCTGAGGTCTCCCCTTCCCTATATGGTTTTAATCCAGCATTCAGTAGATGCAAAGCATGGAAGCAATAAACAGTATCATCCAGGGACAGCCTGGGGT
TGTGTGACACCAGACTCACACCTACACCATCAGTCTCTGAGCAGCATTGCCAGCAAGAGTGGGGCTAGAGGCTTGTCCTTGTCACCCTCGGTCTTCCAGGATTGGTTTTATTGTCCACATTTGTGCAGAATGA
GACACCGAGACCACAGCAGATAGACTCCTCGCTCTAAGAGGCTTGTAGCAGAGTAGAGGCCGATGCTAGACCCCAGAAAGACCTTGAAGCACCATGCAAGGAGCTCAGGACCTAGAGGAGTGTGGGAGT
TGAAAGTTGTCATCGATCCACCCCGTGCAGTGCTCAGTCCCAGAGGGCAGGGTGTGGGGCGCAGGATGACAAAACCCGGAGAGAAATGCAGCGCTGGGTGCGGCCAGGTTATGAAATGGCCTCTGAGC
TCAGCCCCAAATTCTCTTCTTTTCTGCGGTAGGGGCTAGAGGGACGGTAGGTACCCTATGTGGGTATGGAAGCTCCGCCTTCAGCATCCTCAGCCTCCACAGGATATTCAGGGAGCAGCCATGGGGACAGA
AGCAAGGCCTGAATGCAAGGGGTGGAGGGGAGACAGAGAGAGCTGAGGGGAACAGACTATAAGTTCCTTGACCACGGGACCACAGAAAGATTGACAACCCCCACCCCCCAGCATCCTGTTGTATCAGTAT
GACCTAGAGATCACACTAGACTTTCTACAACCAGGTTCAAAGCTTCCAGCTGCGTT

Color legend 5' arm — GFP — P2A — 3' arm
