## Supplementary figures and images for "GFP Reporter System Reveals Cell-to-Cell Variability in Aquaporin-2 Expression"

### Supplemental Figure 2

A

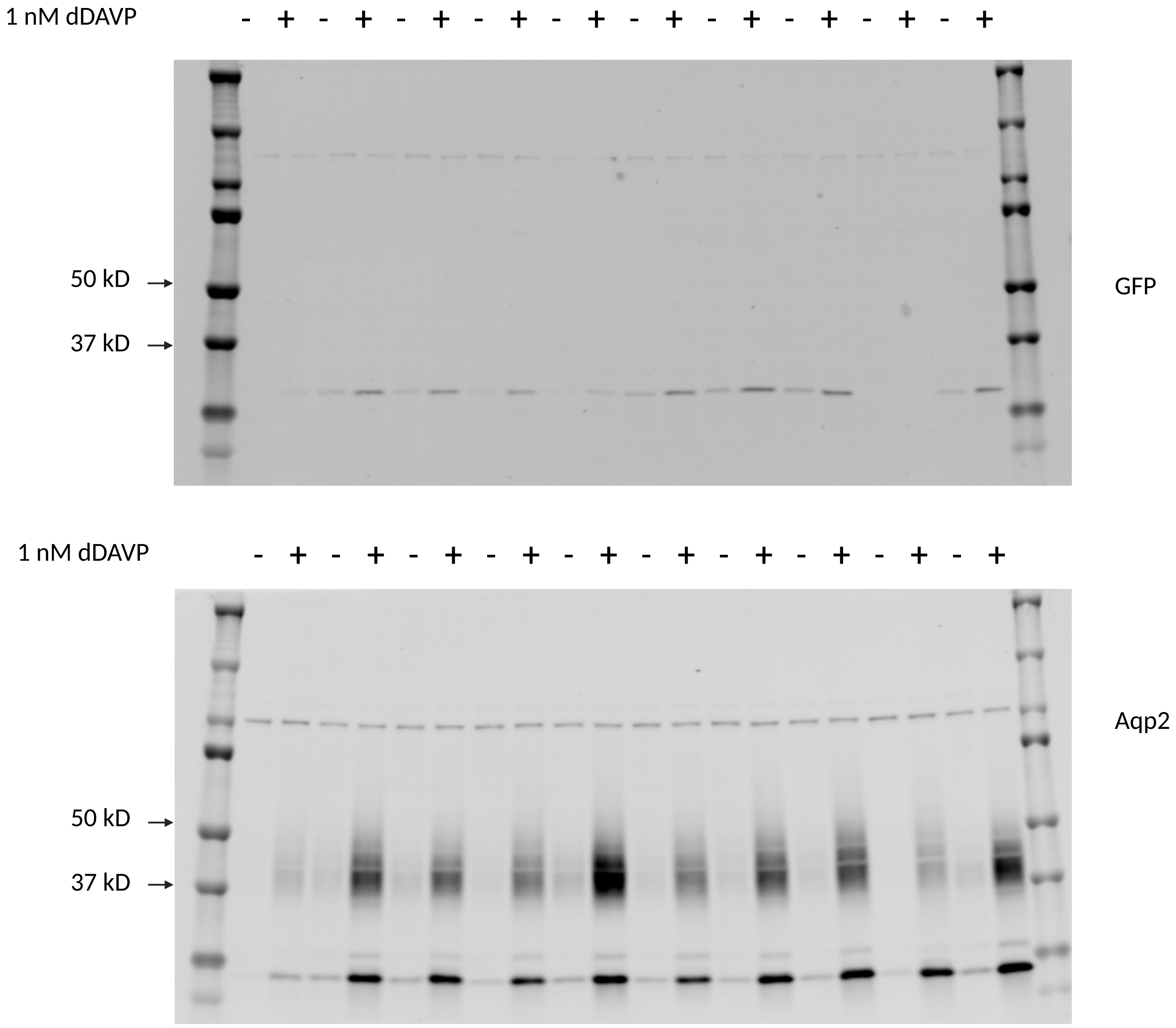
